# From image formation to image representation: the human visual system preserves the hierarchy of 2-dimensional pattern regularity

**DOI:** 10.1101/223743

**Authors:** Peter J. Kohler, Anthony M. Norcia

**Affiliations:** Department of Psychology, Stanford University, Jordan Hall, Building 420, 450 Serra Mall, Stanford, CA 94305, USA

## Abstract

Symmetries are present at many scales in images of natural scenes, due to a complex interplay of physical forces that govern pattern formation in nature. The importance of symmetry for visual perception has been known at least since the gestalt movement of the early 20^th^ century. Since then, symmetry has been shown to contribute to the perception of shapes[1, 2] scenes[3] and surface properties[4], as well as the social process of mate selection[5]. In the two spatial dimensions relevant for images, the four fundamental symmetries, reflection, rotation, translation and glide reflection, can be combined in 17 distinct ways, the “wallpaper” groups[6–8]. The 17 wallpaper obey a hierarchy of complexity, determined by mathematical group theory, where simpler groups are subgroups of more complex ones[9]. Here we use Steady-State Visual Evoked Potentials (SSVEPs) to measure responses to the complete set of wallpaper groups, and present a complete description of the neural basis of symmetry. We find that activity in human visual cortex is remarkably consistent with the hierarchical relationships between the wallpaper groups. Specifically, the amplitudes of symmetry-specific responses in individual participants (n=25) preserve these relationships at an above-chance level in 88.3% (53 out of 60) of cases. Visual cortex thus encodes all the fundamental symmetries using a representational structure that closely approximates the subgroup relationships from group theory. Given that most participants had no knowledge of group theory, the ordered structure of visual responses to wallpaper groups is likely learned implicitly from regularities in the visual environment.

## Results

The visual stimuli for our experiment were multiple exemplar images belonging to each of the 17 wallpaper groups, generated from random-noise textures, as described in detail elsewhere[10]. To isolate brain activity specific to the symmetry structure in the images from activity associated with modulation of low-level features, we used a steady-state design, in which exemplar images belonging to 16 of the 17 wallpaper groups alternated with phase-scrambled images of the same group. Because all wallpapers are periodic images due to their lattice tiling structure, the phase-scrambled images are also a wallpaper group (P1). P1 contains no symmetries other than translation, while all other groups contain translation in combination with one or more of the other three fundamental symmetries (reflection, rotation, glide reflection)[8].

Exemplars from each of the 16 groups alternated at 0.83 Hz with their corresponding set of P1 exemplars, that were matched in terms of their Fourier power spectrum. Because the P1 group serves a control stimulus in this approach, the experiment was restricted to the remaining 16 groups. This design allows us to isolate responses to structural features beyond the shared power spectrum, including any symmetries other than translation, in the odd harmonics of the image update frequency[10–12]. Thus, the magnitude of the odd harmonic response components can be used as a distance metric for each group, with distance being measured relative to the simplest group, P1.

A wallpaper group is a topologically discrete group of isometries of the Euclidean plane, i.e. transformations that preserve distance[8]. Wallpaper groups differ in the number and kind of these transformations. In mathematical group theory, when the elements of one group is completely contained in another, the inner group is called a subgroup of the outer group[13]. Subgroup relationships between wallpaper groups can be distinguished by their indices. The index of a subgroup relationship is the number of cosets, i.e. the number of times the subgroup is found in the outer group[13]. As an example, let us consider groups P6 and P2. If we ignore the translations in two directions that both groups share, group P6 consists of the set of rotations {0°, 60°, 120°, 180°, 240°, 300°}, in which P2 {0°,180°} is contained. P2 is thus a subgroup of P6, and the full P6 set can be generated by every combination of P2 and rotations {0°, 120°, 240°}. Because P2 is repeated three times in P6, P2 is a subgroup of P6 with index 3[14].

The subgroup relationships among the 16 wallpaper groups, as identified by Coxeter & Moser[9] are shown as directed graphs in Figure 1. In a few cases, two groups are subgroups of each other. This is the case for three pairs of groups: PM-CM, PMM-CMM, and P3M1-P31M. These bidirectional relationships were not included in the analysis described below, as was the subgroup relationship that each group has with itself[9]. Note that subgroup relationships with indices of 8 or more (4 in total among the 16 wallpaper groups) were also excluded from the analysis and Figure 1. This resulted in a total number of 60 relationships (shown with connecting arrows in Figure 1). Although the subgroup relationships are not always obvious perceptually, they provide an objective hierarchy among the wallpaper groups that can be compared to the brain data.

**Figure 1:**
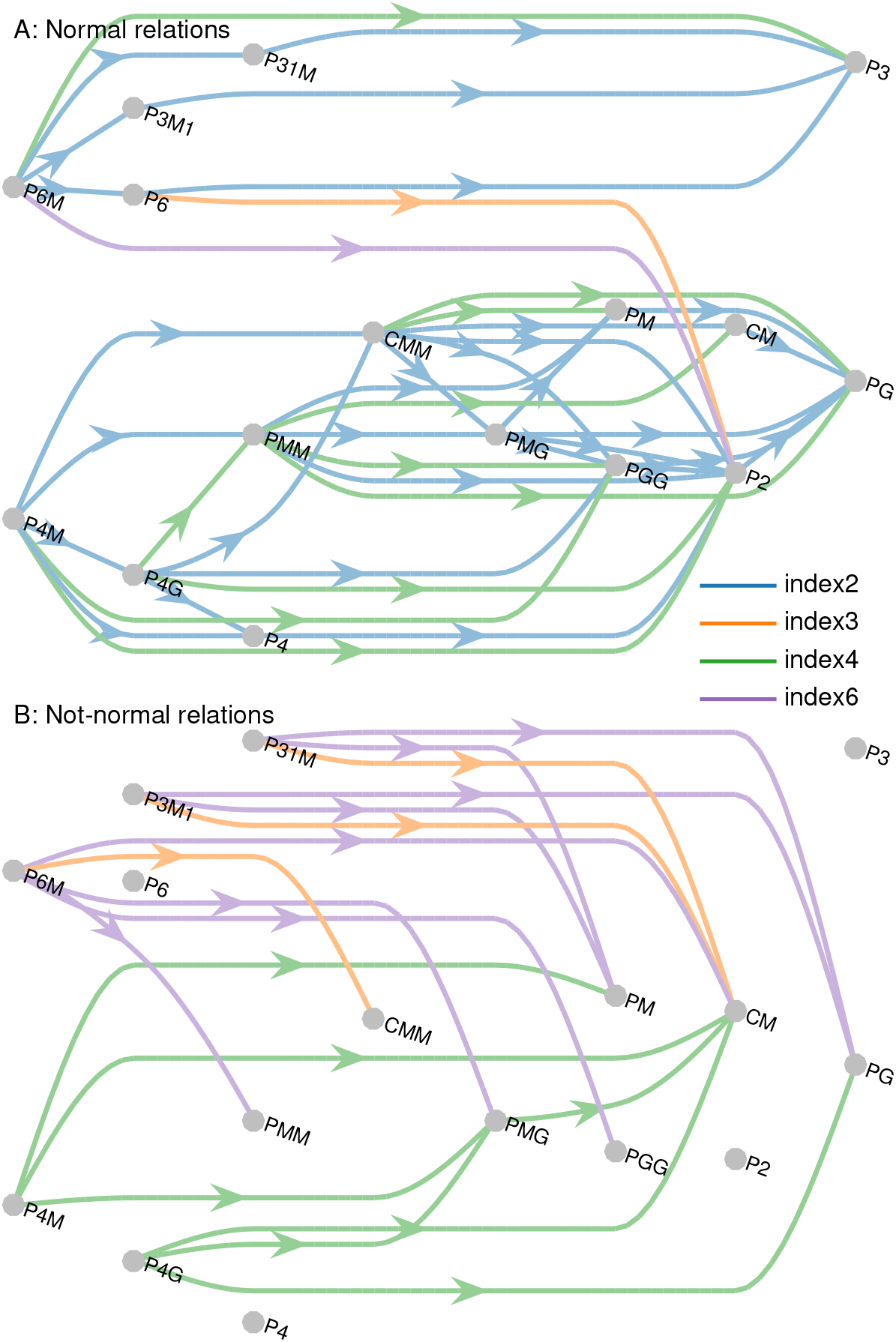
The subgroup relationships between the 16 wallpaper groups studied here. Extracted from Table 4 in Coxeter and Moser[9]. Relationships are shown separately for subgroups that are normal **(A)** and subgroups that are not normal **(B)**. The different colors indicate the index of the subgroup as a subgroup of the outer group. Subgroup relationships with index 2 are always normal. Subgroup relationships with index of 8 or more are very rare, and were therefore not included in the analysis or the figure.

We acquired EEG from 128 electrodes in 25 participants, who each did 16 trials, each with a duration of 12 seconds, of each of the 16 wallpaper groups. This resulted in a total of 256 trials (~51 minutes of data) per participant, which were collected in two separate sessions. After preprocessing, we Fourier-transformed the data and separated the odd and even harmonics, before back-transforming the data into the time-domain. In the following analysis, we focused on the symmetry-specific odd harmonics averaged within a six-electrode region-of-interest (ROI) over occipital cortex, but we also did control analyses using the even harmonics from the occipital ROI and using the odd harmonics from a six-electrode ROI over parietal cortex.

We made contact between the subgroup relationships from group theory and the symmetry-specific brain responses using a straightforward approach. After averaging waveforms over the ROI, we computed the root-mean-square (RMS) over time for each participant’s data. This provided a single number capturing the amplitude of the SSVEP for each group. Odd-harmonic waveforms averaged over participants, for each of the 16 wallpaper groups, are shown in in descending order by RMS magnitude in Figure 2, along with exemplar images that were used in the experiment. For each subgroup relationship, we tested if the RMS was smaller for the subgroup than the outer group, for each of our 25 participants. If this was true for a significant number of participants, we concluded that this subgroup relationship was preserved in the brain data. The RMS metric was thus used as a means of establishing the partial ordering of the neural responses to the wallpaper groups, which could then be compared with the partial ordering based on subgroup relationships from group theory. We determined significance as a cumulative binomial probability, reasoning that for each participant the RMS could either be smaller for the subgroup or not, given us 50% chance of success. This resulted in a threshold for significance of 18 or more participants out of 25 (*p* < 0.022), but we also applied a stricter criterion of 20 or more participants (*p* < 0.002).

**Figure 2:**
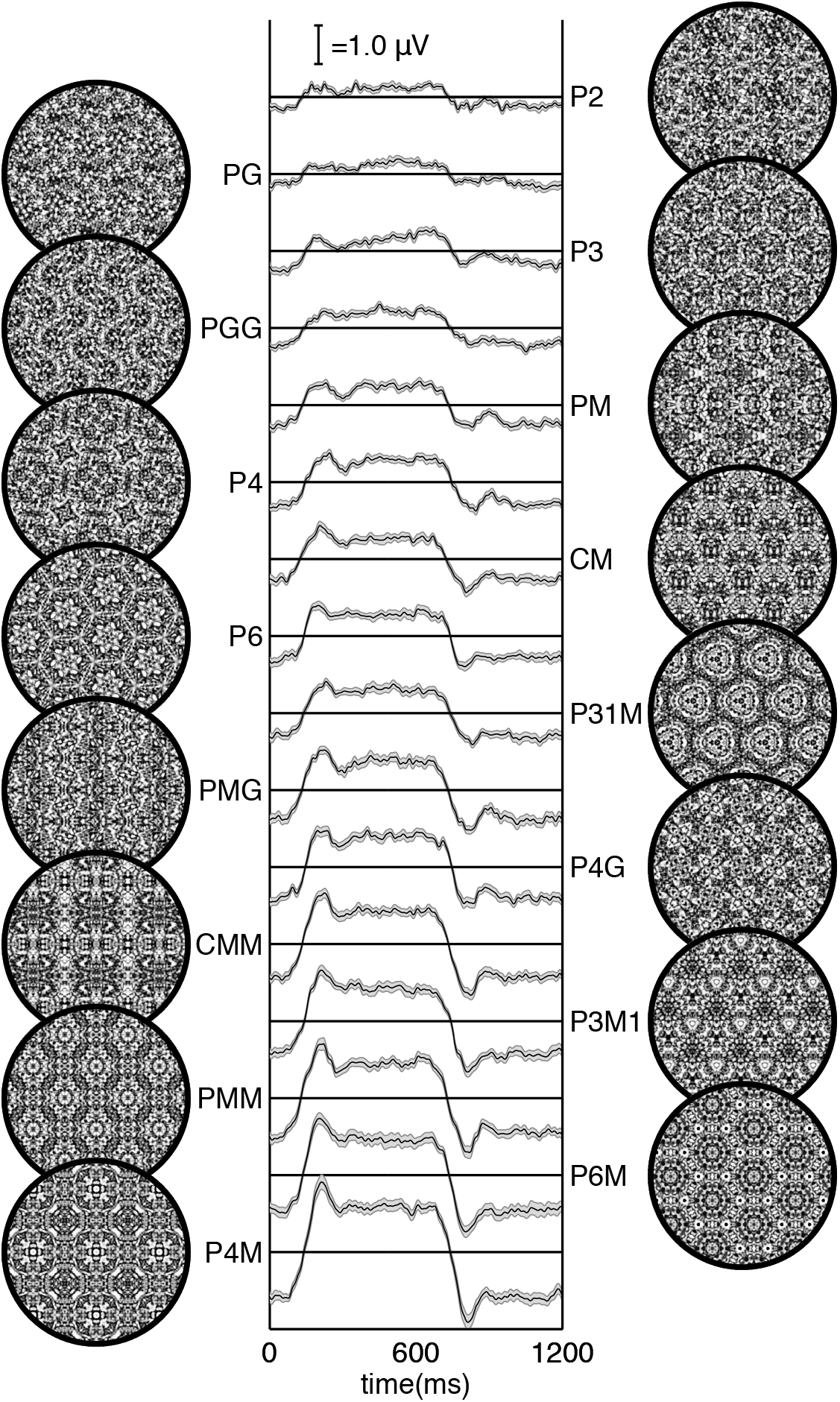
Steady-Visual Evoked Potentials to the 16 wallpaper groups. Generated by isolating the odd harmonics of the evoked response waveforms, averaged across a six-electrode region-of-interest (ROI) over occipital cortex. The waveforms represent the differential response between each of the groups shown and the P1 group, which contained no symmetries besides translation. Example images from each of the 16 groups are shown next to each waveform. The waveforms are arranged in descending order by mean RMS score across participants. Supplemental Figure S1 shows the location on the scalp of the six-electrode occipital ROI, as well as the location of a second six-electrode ROI over parietal cortex. Supplemental Figure S2 shows analogous waveforms in which the even harmonics have been isolated, and Supplemental Figure S3 shows the odd harmonics from the parietal cortex ROI. While the odd harmonics over occipital cortex display differ strongly in their amplitude for each group, this is much less evident in the even harmonics, and in the odd harmonics over parietal cortex.

The brain data preserved 53 out of 60 subgroup relationships in a significant number of participants at the liberal threshold (88.3 %; darker dotted line in Figure 3). 49 out of 60 relationships (81.6%) were preserved the brain data of enough participants to reach the stricter criterion for significance (lighter dotted line in Figure 3). To provide a measure of how consistently the subgroup relationships were preserved in individual participants, we can report the average number of participants for whom the relationships were preserved, across a set of relationships. For the set of all 60 subgroup relationships, the relationships were preserved in 21.6 out of 25 participants on average. We conclude that most subgroup relationship are preserved in the brain data of most participants.

**Figure 3:**
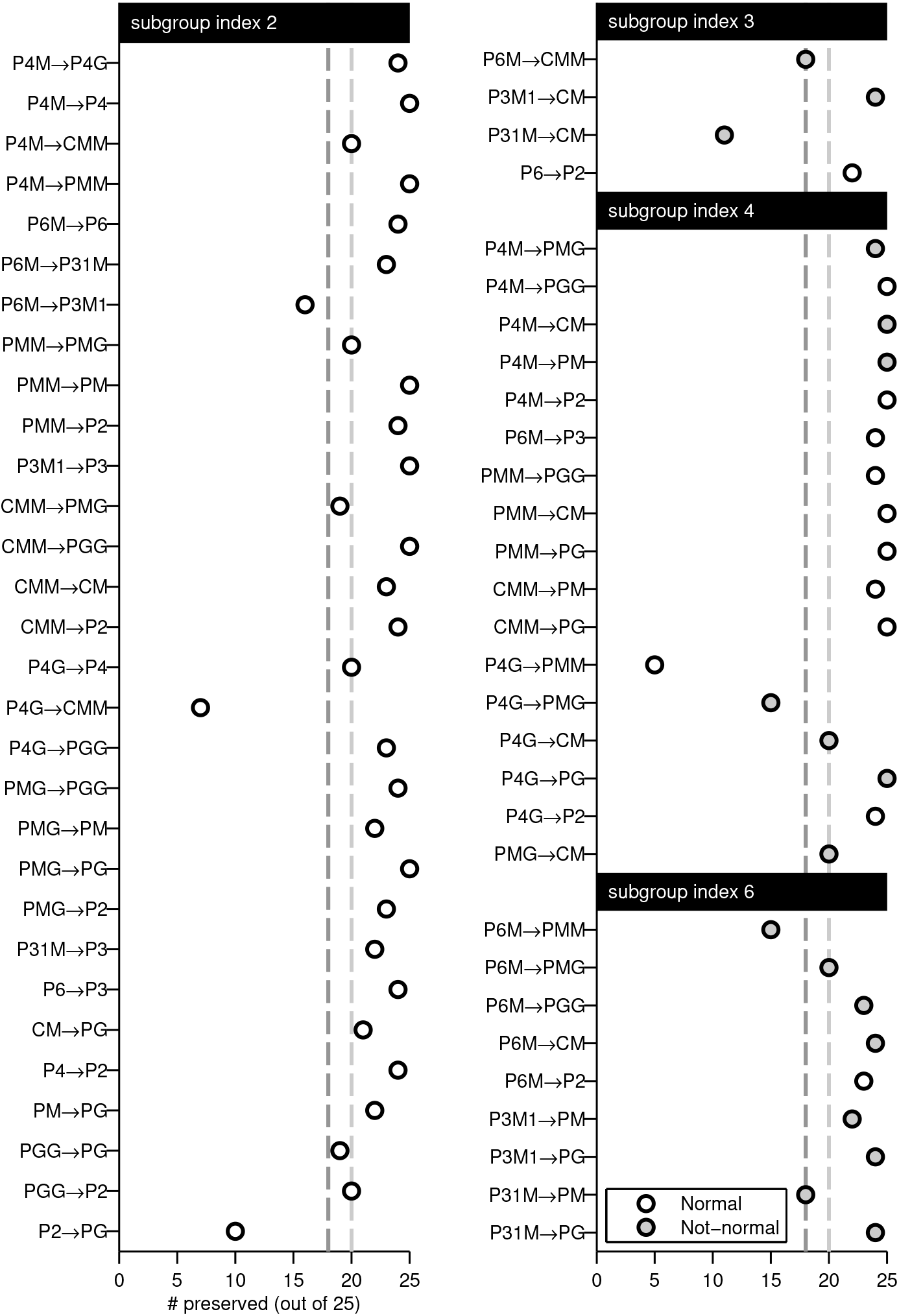
Preservation rates across 60 subgroup relationships, odd harmonics over occipital cortex. Number of participants for which a given subgroup relationship was preserved in the odd harmonics in the occipital cortex ROI, shown separately for each subgroup relationship. Dark dotted line represents the threshold for *p* < 0.022, while light gray dotted line represents the threshold for *p* < 0.002. Subgroup relationships with different indices are plotted separately, and for each index, the relationships are organized by outer group. Open dots are normal subgroup relationships, filled dots are not-normal subgroup relationships. Figure S4 shows the analogous plot for the even harmonics in the same ROI, and Supplemental Figure S5 shows the analogous plot for the odd harmonics from the parietal cortex ROI.

The correspondence between brain data and subgroup relationships was specific to the odd harmonics in occipital cortex. For the even harmonics in the occipital ROI, only 18 out of 60 (30 %) relationships were preserved in a significant number of participants (at *p* < 0.022; see Supplemental Figure 4). For the odd harmonics in the parietal ROI, only 8 out 60 (13.3 %) relationships were preserved in a significant number of participants (at *p* < 0.022; see Supplemental Figure 5). Furthermore, the even harmonics and parietal ROI did not add any information: Any relationship that was significant in the even harmonics over occipital cortex or in the odd harmonics over parietal cortex, was also significant in the odd harmonics over visual cortex. This confirms that our experimental design can isolate symmetry-specific responses in the odd harmonics and shows that the hierarchical structure of the subgroup relationships is preserved by visual cortex.

Subgroup relationships can be either normal and not-normal[14] (shown separately in Figure 1). Normal and not-normal subgroup relationships appeared to be preserved about equally well, with significance rates of 90.4 % (38 of 42) and 83.3 % (15 of 18), respectively. Normal subgroups were preserved in 21.8 participants on average, while not-normal were preserved in 20.9 participants on average. The different subgroup indices also lead to similar preservation rates, with perhaps slightly worse performance for index 3 (see Figure 3).

Are some wallpaper groups better characterized as outer groups or subgroups? The outer group that produced highest preservation rate was P4M, for which all 9 relationships were preserved in a significant number of participants (average number of participants: 24.2). The outer group that produced lowest preservation was P2, whose single relationship (with subgroup PG) was only preserved in 10 participants, not enough to reach significance. P4G also had relatively low preservation, only 4 of its 6 outer group relationships reached significance (average number of participants: 17.2). All relationships in which PGG (6 of 6) or P3 (3 of 3) were subgroups were significantly preserved, in 23.8 participants on average (P6 and P4G also had high preservation, 24 of 25 participants, but only one relationships in which each was the subgroup). PMM was the subgroup that produced the poorest preservation rates, with 1 of 3 relationships reaching significance, and an average number of participants of 15.

## Discussion

Here we show that beyond merely responding to the elementary symmetry operations of reflection[15] and rotation[10], the visual system explicitly represents hierarchical structure of the 17 wallpaper groups, and thus the compositions of all four of the fundamental symmetry transformations (rotation, reflection, translation, glide reflection) which comprise regular textures. The RMS measure of SSVEP amplitude, preserves the complex hierarchy of subgroup relationships among the wallpaper groups[9]. Out of a total of 60 relationships, 53 were preserved in a significant number of participants, and 49 were significant even at a stricter threshold (*p* < 0.002). The ordering was highly stable in individual participants, with an average preservation rate of ~21 of 25 participants across all 60 relationships (see Figure 3).

This remarkable consistency was specific to the odd harmonics of the stimulus frequency, that capture the symmetry-specific response[10] and to electrodes in an ROI over occipital cortex. When the same analysis was done on the even harmonics of the occipital cortex ROI, the ordering of responses was much less apparent (see Figure S2) and preservation rates much lower (see Figure S4). The odd harmonics from electrodes in an ROI over parietal cortex, showed even weaker evidence of preserving the hierarchy among sub-groups (see Figure S5). Importantly, no relationships were preserved in either of these control analyses that were not also preserved in the main analysis of the odd harmonics in the occipital cortex ROI.

The current data provide a complete description of the visual system’s response to symmetries in the 2-D plane. Our design does not allow us to independently measure the response to P1, but because each of the 16 other groups produce non-zero odd harmonic amplitudes (see Figure 2), we can conclude that the relationships between P1 and all other groups, where P1 is the subgroup, are also preserved by the visual system.

The subgroup relationships are not obvious perceptually, and most participants had no knowledge of group theory. Thus, the visual system’s ability to preserve the subgroup hierarchy does not depend on explicit knowledge of the relationships. Furthermore, behavioral experiments have shown that although naïve observers can distinguish many of the wallpaper groups[16], they are generally error-prone when asked to assign exemplar images to the appropriate group[17].

The correspondence between responses in the visual system and group theory that we demonstrate here, may reflect a form of implicit learning that depends on the structure of the natural world. The environment is itself constrained by physical forces underlying pattern formation and these forces are subject to multiple symmetry constraints[18]. The ordered structure of responses to wallpaper groups could be driven by a central tenet of neural coding, that of efficiency. If coding is to be efficient, neural resources should be distributed in such a way that the structure of the environment is captured with minimum redundancy considering the visual geometric optics, the capabilities of the subsequent neural coding stages and the behavioral goals of the organism[19–22]. Early work within the efficient coding framework suggested that natural images had a 1/f spectrum and that the corresponding redundancy between pixels in natural images could be coded efficiently with a sparse set of oriented filter responses, such as those present in the early visual pathway[23, 24]. Our results suggest that the principle of efficient coding extends to a much higher level of structural redundancy – that of symmetries in visual images.

The 17 wallpaper groups are completely regular, and relatively rare in the visual environment, especially when considering distortions due to perspective and occlusion. Near-regular textures, however abound in the visual world, and can be approximated as deformed versions of the wallpaper groups[25]. The correspondence between brain data and group theory demonstrated here may indicate that the visual system represents visual textures using a similar scheme, with the wallpaper groups serving as anchor points in representational space. This framework resembles norm-based encoding strategies that have been proposed for other stimulus classes, most notably faces[26], and leads to the prediction that adaptation to wallpaper patterns should distort perception of near-regular textures, similar to the aftereffects found for faces[27].

Field biologist have demonstrated that animals respond more strongly to exaggerated versions of a learned stimulus, referred to as “supernormal” stimuli[28]. In the norm-based encoding framework, wallpaper groups can be considered super-textures, exaggerated examples of the near-regular textures that surround us. Artists may consciously or unconsciously create supernormal stimuli, to capture the essence of the subject and evoke strong responses in the audience[29]. Wallpaper groups are visually compelling and have been widely used in human artistic expression going back to the Neolithic age[30]. If wallpapers are super-textures, their prevalence may be a direct consequence of the strategy the human visual system uses for encoding visual textures.

## Acknowledgments

This research was supported by a NSF grant INSPIRE 248076. The authors thank Yanxi Liu for helpful discussions about wallpaper groups and group theory.

## Author Contributions

Conceptualization, P.J.K. and A.M.N.; Methodology, P.J.K. and A.M.N.; Investigation, P.J.K., Formal Analysis, P.J.K.; Visualization, P.J.K.; Writing, P.J.K. and A.M.N.; Funding Acquisition, A.M.N.; Resources, A.M.N.

## References

1. Palmer, S.E. (1985). The role of symmetry in shape perception. Acta psychologica 59, 67–90.

2. Li, Y., Sawada, T., Shi, Y., Steinman, R., and Pizlo, Z. (2013). Symmetry Is the sine qua non of Shape. In Shape Perception in Human and Computer Vision, S.J. Dickinson and Z. Pizlo, eds. (Springer London), pp. 21–40.

3. Apthorp, D., and Bell, J. (2015). Symmetry is less than meets the eye. Current Biology 25, R267–R268.

4. Cohen, E.H., and Zaidi, Q. (2013). Symmetry in context: Salience of mirror symmetry in natural patterns. Journal of vision 13.

5. Møller, A.P. (1992). Female swallow preference for symmetrical male sexual ornaments. Nature 357, 238–240.

6. Fedorov, E. (1891). Symmetry in the plane. In Zapiski Imperatorskogo S. Peterburgskogo Mineralogichesgo Obshchestva [Proc. S. Peterb. Mineral. Soc.], Volume 2. pp. 345–390.

7. Polya, G. (1924). XII. Über die Analogie der Kristallsymmetrie in der Ebene. Zeitschrift für Kristallographie-Crystalline Materials 60, 278–282.

8. Liu, Y., Hel-Or, H., Kaplan, C.S., and Van Gool, L. (2010). Computational Symmetry in Computer Vision and Computer Graphics. Foundations and Trends^®^ in Computer Graphics and Vision 5, 1–195.

9. Coxeter, H.S.M., and Moser, W.O.J. (1972). Generators and relations for discrete groups, (Berlin, New York: Springer-Verlag).

10. Kohler, P.J., Clarke, A., Yakovleva, A., Liu, Y., and Norcia, A.M. (2016). Representation of Maximally Regular Textures in Human Visual Cortex. The Journal of Neuroscience 36, 714–729.

11. Norcia, A.M., Appelbaum, L.G., Ales, J.M., Cottereau, B.R., and Rossion, B. (2015). The steady-state visual evoked potential in vision research: A review. Journal of vision 15, 4–4.

12. Norcia, A.M., Candy, T.R., Pettet, M.W., Vildavski, V.Y., and Tyler, C.W. (2002). Temporal dynamics of the human response to symmetry. Journal of vision 2.

13. Carter, N.C. (2009). Visual group theory.

14. Carter, N.C. (2009). Visual Group Theory, (Washington, UNITED STATES: Mathematical Association of America).

15. Sasaki, Y., Vanduffel, W., Knutsen, T., Tyler, C., and Tootell, R. (2005). Symmetry activates extrastriate visual cortex in human and nonhuman primates. Proceedings of the National Academy of Sciences of the United States of America 102, 3159–3163.

16. Landwehr, K. (2009). Camouflaged symmetry. Perception 38, 1712–1720.

17. Clarke, A.D.F., Green, P.R., Halley, F., and Chantler, M.J. (2011). Similar Symmetries: The Role of Wallpaper Groups in Perceptual Texture Similarity. Symmetry 3, 246–264.

18. Hoyle, R.B. (2006). Pattern formation: an introduction to methods, (Cambridge University Press).

19. Laughlin, S. (1981). A simple coding procedure enhances a neuron's information capacity. Zeitschrift fur Naturforschung. Section C: Biosciences 36, 910–912.

20. Attneave, F. (1954). Some informational aspects of visual perception. Psychological review 61, 183–193.

21. Barlow, H.B. (1961). Possible principles underlying the transformations of sensory messages. In Sensory Communication, W.A. Rosenblith, ed. (MIT Press), pp. 217–234.

22. Geisler, W.S., Najemnik, J., and Ing, A.D. (2009). Optimal stimulus encoders for natural tasks. Journal of vision 9, 17–17.

23. Field, D.J. (1987). Relations between the statistics of natural images and the response properties of cortical cells. Journal of the Optical Society of America. A, Optics and image science 4, 2379–2394.

24. Olshausen, B.A., and Field, D.J. (1997). Sparse coding with an overcomplete basis set: a strategy employed by V1? Vision research 37, 3311–3325.

25. Liu, Y., Lin, W.-C., and Hays, J. (2004). Near-regular texture analysis and manipulation. In ACM Transactions on Graphics (TOG), Volume 23. (ACM), pp. 368–376.

26. Leopold, D.A., Bondar, I.V., and Giese, M.A. (2006). Norm-based face encoding by single neurons in the monkey inferotemporal cortex. Nature 442, 572–575.

27. Webster, M.A., and MacLin, O.H. (1999). Figural aftereffects in the perception of faces. Psychon Bull Rev 6, 647–653.

28. Tinbergen, N. (1953). The herring gull's world: a study of the social behaviour of birds, (Oxford, England: Frederick A. Praeger, Inc.).

29. Ramachandran, V.S., and Hirstein, W. (1999). The science of art: A neurological theory of aesthetic experience. Journal of Consciousness Studies 6, 15–41.

30. Jablan, S.V. (2014). Symmetry, Ornament and Modularity, (Singapore, SINGAPORE: World Scientific Publishing Co Pte Ltd).

